# Altered coding of environmental boundaries in human aging: an fMRI study

**DOI:** 10.1101/2025.09.25.678564

**Authors:** Vladislava Segen, Matthias Stangl, Jonathan Shine, Thomas Wolbers

**Affiliations:** Aging, Cognition & Technology Group, German Center for Neurodegenerative Diseases (DZNE), Magdeburg 39120, Germany; Department of Biomedical Engineering and Department of Psychological and Brain Sciences, Center for Systems Neuroscience, Cognitive Neuroimaging Center, Neurophotonics Center, Boston University, Boston, MA 02215, USA; Department of Neurosurgery, Boston Medical Center, Boston University Chobanian and Avedisian School of Medicine, Boston, MA 02118, USA; Heartbeat Medical, Berlin, 10829, Germany; Center for Behavioural Brain Sciences (CBBS), Otto-von-Guericke-University Magdeburg, Magdeburg 39120, Germany

**Author notes:** Joint senior authors (Jonathan Shine, Thomas Wolbers). Corresponding author: Vladislava Segen.

## Abstract

Aging is associated with changes in spatial memory and navigation, yet the mechanisms underlying these changes are not yet fully understood. Environmental boundaries are among the most salient and reliable spatial cues, supporting both spatial memory and orientation. Here, we investigated how aging affects the use and the neural representation of boundary information during a virtual object location memory task. Healthy young and older adults navigated a square virtual environment while undergoing functional magnetic resonance imaging, allowing us to assess moment-to-moment encoding of distance to environmental boundaries in the entorhinal cortex and subiculum. Behaviorally, both age groups showed more accurate memory for objects located near boundaries, but this effect was amplified in older adults, whose spatial precision declined more steeply with increasing distance from boundaries. Older adults also exhibited a stronger bias to recall objects closer to boundaries. Analysis of navigation behaviour revealed that older adults followed boundary-oriented paths regardless of target location, whereas young adults flexibly adapted their navigation based on spatial context. Neurally, older adults—but not young adults—showed significant blood-oxygen-level-dependent modulation by boundary distance in the entorhinal cortex and subiculum, with activity decreasing as participants moved farther from boundaries. This effect was most pronounced in low-performing older adults and was associated with stronger behavioral boundary bias, suggesting a maladaptive reliance on proximity-based cues. Together, our results provide converging behavioral and neural evidence that aging alters the use and representation of boundary information, with downstream effects on spatial memory. Altered boundary processing may represent a key mechanism contributing to age-related declines in spatial cognition.

## 1 INTRODUCTION

Aging is associated with deficits in spatial navigation and memory (Lester et al., 2017). In healthy aging, these deficits often manifest as a decline in the accuracy and precision of spatial representations (McAvan et al., 2021; Moffat & Resnick, 2002; Segen et al., 2021), likely reflecting changes in the neural systems that support spatial cognition (Barnes, 1988; Diersch et al., 2021; Lester et al., 2017; Stangl et al., 2018). One of the most salient and stable features of natural environments - boundaries - provide critical cues that support spatial orientation and memory. Environmental boundaries, such as walls or edges of paths, help anchor spatial representations and are known to influence memory for object locations (Brunec et al., 2018; Lee et al., 2018). At the neural level, boundary-related coding is supported by specialized cells in the entorhinal cortex (EC) and subiculum, including border cells (Bjerknes et al., 2014; Solstad et al., 2008) and boundary vector cells (Lever et al., 2009) which respond to proximity to environmental edges. In rodents, hippocampal place cells are also modulated by environmental geometry and boundary structure, with place fields shifting in response to changes in boundary layout (O’Keefe & Burgess, 1996), likely reflecting input from boundary-sensitive cells in EC and subiculum. These spatial firing patterns are directly linked to memory performance (O’Keefe & Speakman, 1987), suggesting that hippocampus-dependent memory is influenced by boundary cues (Bicanski & Burgess, 2018; Hartley et al., 2000). In humans, evidence of boundary-related processing has been primarily found across the hippocampus, EC and subiculum (Doeller et al., 2008; Lee et al., 2018; Shine et al., 2019; Stangl et al., 2021), further supporting the importance of this system in anchoring spatial memory.

How does human aging affect the processing of environmental boundaries? Bécu et al. (2020) demonstrated that healthy older adults rely more heavily on geometric cues during real-world navigation. In their study, older adults preferred to use room geometry to navigate within a rectangular environment, even when both landmarks and boundaries were available, suggesting preserved or even enhanced use of structural features. In contrast, Schuck et al. (2015) used a functional magnetic resonance imaging (fMRI) object location memory task in a circular virtual environment and found that older adults were less sensitive to boundary manipulations—specifically changes in arena size—compared to young adults. Young adults adjusted their spatial estimates accordingly and showed a positive association between hippocampal activity and boundary sensitivity, in line with models positing that hippocampal place cells are influenced by boundary vector cell input (Bicanski & Burgess, 2018; Hartley et al., 2000; Lever et al., 2009). This hippocampal–boundary sensitivity relationship was absent in older adults, suggesting an age-related decoupling between boundary-related signals and hippocampal spatial coding. Importantly, these findings were replicated in an independent sample (Baeuchl et al., 2023).

Schuck et al. (2015) provided early insights, showing that older adults were less sensitive to boundary manipulations and exhibited reduced hippocampal responses during spatial memory retrieval. However, there has been little further investigation into how boundary processing is altered in aging, particularly at the neural level and during active navigation. As such, it remains an open question whether age-related changes in boundary use are accompanied by corresponding changes in neural representations in medial temporal lobe structures, and whether these changes reflect compensatory adaptations or a more fundamental disruption of spatial coding mechanisms.

To gain further insight into boundary use during navigation in aging, we re-analyzed data from a self-guided virtual object location memory task (Stangl et al., 2018), in which both healthy younger and older adults learned the locations of objects placed at varying distances from environmental boundaries. The task was designed to mimic navigation, allowing us to capture participants’ navigation behavior and response patterns. Crucially, this approach enabled us to examine boundary processing on a moment-to-moment basis, rather than during isolated recall or feedback phases, which have been the focus of prior work (e.g. Baeuchl et al., 2023; Schuck et al., 2015). This distinction is critical, as spatial computations related to boundary proximity likely unfold continuously during movement and decision-making. We combined behavioral (response and dynamic navigation), and fMRI data to investigate how boundaries are used and encoded during active navigation in healthy young and older adults. Our neural analyses focused on the EC and subiculum, regions which support dynamic boundary-based spatial coding (Lee et al., 2018; Lever et al., 2009; Shine et al., 2019; Solstad et al., 2008; Stangl et al., 2021). By linking real-time boundary distance to Blood-Oxygenation-Level Dependent (BOLD) activity, spatial responses, and navigational behavior, we aimed to clarify how boundary information is dynamically represented during movement in a virtual environment, and whether its role in spatial memory changes with age.

## 2 MATERIALS AND METHODS

### 2.1 Participants

Twenty young adults (M = 24.5, SD = 3.3 years; 10 females, 10 males; age range = 19–30 years) and 21 older adults (M = 69.3, SD = 4.8 years; 11 females, 10 males; age range = 63–81 years) took part in the study. All participants were right-handed, reported normal or corrected-to-normal vision, and had no history of neurological or psychiatric disorders, nor any motor impairments affecting walking or standing. Prior to participation, all individuals completed the Montreal Cognitive Assessment (MoCA) as a screening tool for mild cognitive impairment (Nasreddine et al., 2005). Only participants who scored above the cut-off of 23 (Carson et al., 2018) were included in the study. All participants provided written informed consent, and the study was approved by the Ethics Committee of the University of Magdeburg.

### 2.2 Experiment Paradigm and procedure

The original paradigm has been described in detail elsewhere (Stangl et al., 2018); the present study involves a reanalysis of the dataset published in that work. Briefly, the paradigm consisted of an object location memory task in which participants were required to recall the location of three different target objects (ball, plant and trash bin) in a square virtual environment measuring 160*160 virtual metres (vm). Critically, the objects were positioned at varying distances from the boundaries (Figure 1), allowing for an examination of how proximity to environmental boundaries influences spatial memory performance. Specifically, the total distance to the two closest boundaries was 49 vm for the trash bin (x = 33 vm, y = 16 vm), 96 vm for the plant (x = 27 vm, y = 69 vm), and 102 vm for the ball (x = 64 vm, y = 38 vm). While rodent studies typically use arenas much smaller relative to body size, boundary information remains perceptually meaningful at human scales. Distance estimation in large environments relies on visual cues: as boundaries become closer, they occupy more of the visual field (angular size), and texture gradients on the floor and walls provide reliable depth information and scene layout (Gibson, 1946; Howard & Rogers, 2002; Renner et al., 2013). Our VR environment was designed with continuous floor and wall textures as well as wall-mounted objects, thereby providing rich visual cue input comparable to natural conditions and ensuring that boundary distance remained perceptually accessible even at these scales (Renner et al., 2013).

**Figure 1.**
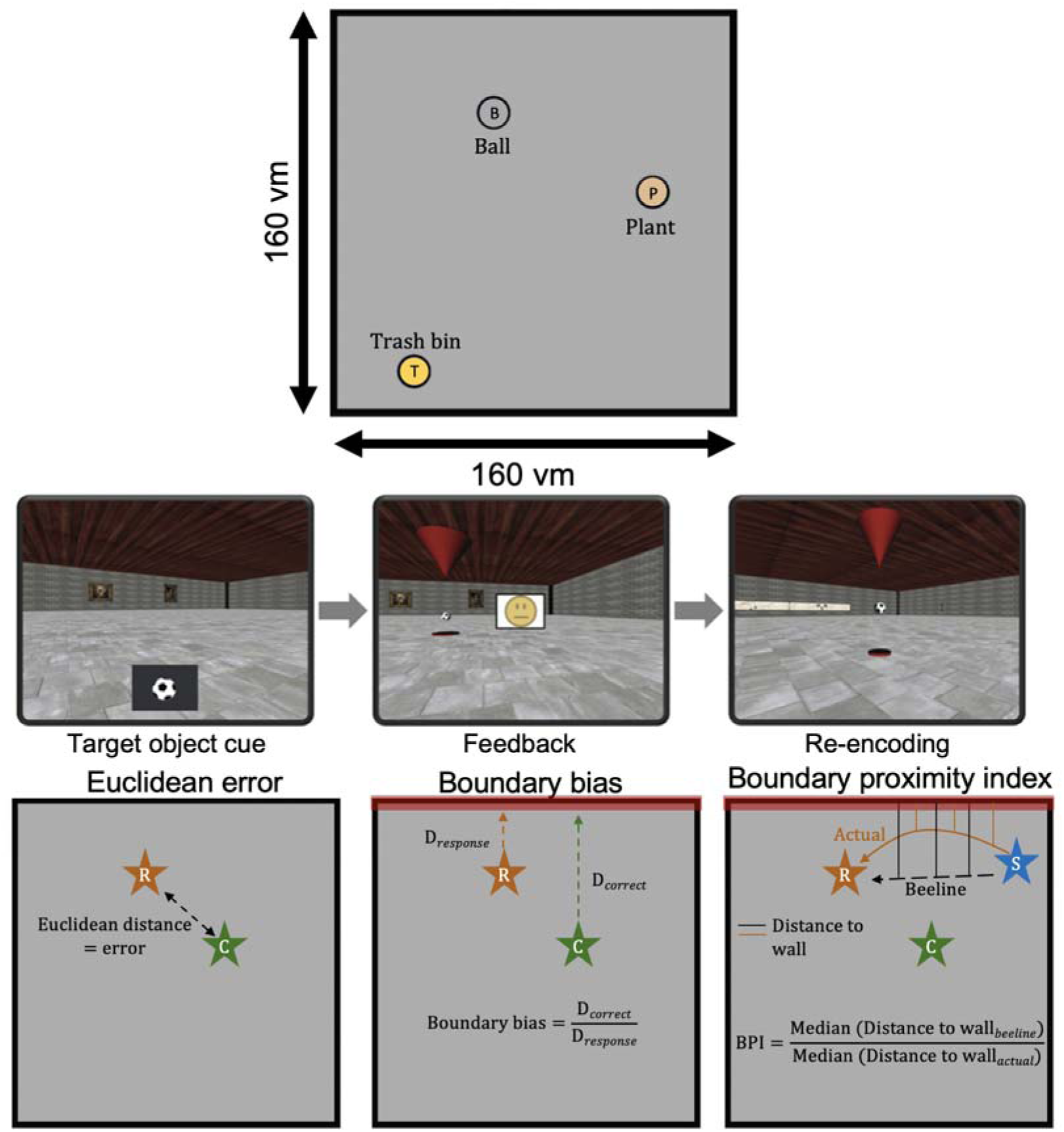
Overview of object location memory task and key behavioral measures. (Top) Participants navigated within a 160 virtual meter (vm) square environment to memorize and recall the locations of three objects (ball, plant, trash bin). (Middle) During each trial, participants were first shown a target object cue, then navigated to the remembered location (left), received feedback (middle), and in case their responses were outside the 20vm range of the target, they were given the opportunity to re-encode the correct location (right). (Bottom) Key behavioral measures included: (1) *Euclidean error*, defined as the straight-line distance between the participant’s response and the correct target location; (2) *Boundary bias*, quantifying the ratio between the distance to the nearest wall between the correct and remembered location; and (3) *Boundary proximity index (BPI)*, which captures navigation bias by comparing the median distance to the nearest wall during actual navigation versus the optimal (beeline) trajectory to the response location. Here, R (orange star) indicates the participant’s response location, C (green star) the correct target location, and S (blue star) the start position at the beginning of each trial.

Each trial followed the same format. The participant was transported to a random location in the virtual environment and a cue comprising one of the three target items would appear on screen. The participant was required to navigate to its remembered location and press a button to confirm their response location. A feedback smiley-face after every trial reflected the participant’s accuracy, with Euclidean distance errors below 20vm eliciting a green smiley; distance errors between 20-30vm elicited a yellow smiley; and distance errors over 30vm resulted in a red smiley. If the error distance was over 20vm the target object was presented again and the participant required to collect it to allow them to re-encode the object location. Linear translation speed was kept constant at 15vm/s and rotation speed was set to 50 deg/s; participants could not perform linear translations whilst rotating meaning that all trajectories comprised straight lines. The height of the virtual camera was set to 1.7vm. The paradigm was developed using WorldViz Vizard 5.1 Virtual Reality Software (WorldViz LLC, http://www.worldviz.com) and participants used an MR compatible joystick (Tethyx, Current Designs, http://www.curdes.com) to move through the virtual environment.

One day prior to scanning, participants were trained extensively on the task. Participants progressed to the scanned test phase once they identified each target object’s location at least two times in a row with an error distance below 20vm, and had received at least eight green smiley faces in a row. To ensure that the participants still remembered the object location associations on the day of scanning, the training task was repeated but this time with the requirement to identify the location of the target items (error < 20vm) on five consecutive trials. Then, each participant completed four runs of fMRI scanning while performing the object location memory task. Each fMRI scanning run had a duration of 16 minutes. Given that we matched the scanning duration, there were differences in the number of trials performed by younger (M = 193.65, SD=16.48) and older adults (M=149.19, SD=15.41).

### 2.3. (f)MRI Acquisition

MRI data were acquired on a 3T Siemens Magnetom Prisma scanner equipped with a 64-channel phased array head coil. All sequences described below utilized parallel imaging with a GRAPPA acceleration factor of 2. A structural T1-weighted image with isotropic resolution was acquired using an MPRAGE sequence with the following parameters: TR = 2500 ms, TE = 2.82 ms, inversion time (TI) = 1100 ms, slice thickness = 1 mm, in-plane-resolution = 1 × 1 mm, number of slices = 192, field of view = 256 mm, flip angle = 7°. For segmentation of the EC and the subiculum, a high-resolution T2-weighted structural image was acquired using a hyper echo turbo-spin-echo (TSE) sequence with the following parameters: TR = 6000 ms, TE = 71 ms, slice thickness = 2 mm, in-plane-resolution = 0.5 × 0.5 mm, number of slices = 64, field of view = 224 mm, flip angle = 120°, slice acquisition order = interleaved. Slices were oriented orthogonal to the long axis of the hippocampus.

During the object location memory task, T2_∗_-weighted functional images were recorded with a partial volume echo-planar imaging (EPI) sequence with the following parameters: repetition time (TR) = 1500 ms, echo time (TE) = 30 ms, slice thickness = 2 mm, in-plane-resolution = 2 × 2 mm, number of slices = 24, field of view = 216 mm, flip angle = 80°, slice acquisition order = interleaved. Slices were oriented parallel to the long axis of the hippocampus. To facilitate an accurate co-registration of EC masks to partial volume EPI images, a whole brain EPI image was acquired with the following parameters: TR = 6000 ms, TE = 30 ms, slice thickness = 2 mm, in-plane-resolution = 2 × 2 mm, number of slices = 84, field of view = 216 mm, flip angle = 90°, slice acquisition order = interleaved. Slices were oriented parallel to the long axis of the hippocampus.

### 2.4. Region of interest mask segmentation and registration

Anatomical masks of the EC were traced manually on each participant’s T2-weighted image using ITK-SNAP (http://www.itksnap.org/), following the segmentation protocol of Berron et al. (2017) Subiculum were automatically segmented using ASHS-T2 atlas (Yushkevich et al., 2015).

Together with the T2-weighted image, ROI mask images were co-registered to the participant’s EPI data. Given that the EPI images were only partial volume slabs, the process of co-registering T2 images to the partial volume EPI images is non-trivial, as co-registration can only be computed on a relatively small portion of overlapping brain tissue between the two different image modalities. Co-registration was therefore performed via one whole brain EPI image, in order to avoid co-registering images with little overlap of brain tissue, and consequently allow for a more accurate co-registration. This whole brain EPI image had similar imaging parameters as the partial volume EPI images, but a considerably higher TR to enable whole brain coverage. Co-registration of the T2 image to the partial volume EPI images included two separate steps: In a first step, the T2 image (together with the ROI masks) was co-registered to the whole brain EPI image. This co-registration step has the advantage that co-registration is facilitated by considerably more overlapping brain tissue between the two image modalities (i.e., T2 and whole brain EPI), as compared to directly co-registering the T2 image to the partial volume EPI images. In a second step, the whole brain EPI (together with the T2 image and ROI masks) was co-registered to the partial volume EPI images. This second co-registration step again has the advantage of considerably more overlap between whole brain EPI and partial volume EPI images. Furthermore, whole brain EPI and partial volume EPI images share similar imaging parameters and therefore have widely similar properties, which also facilitates a more accurate co-registration.

### 2.5 fMRI data processing and analysis

Functional images were realigned and smoothed using a 5 mm FWHM Gaussian kernel in SPM12 (http://www.fil.ion.ucl.ac.uk/spm/). To avoid spatial distortions and interpolation errors from template normalization, all analyses were conducted in each participant’s native space. Standard head motion correction was applied to account for linear and angular displacement. Physiological noise was estimated using anatomical CompCor (aCompCor), which extracts noise components from white matter (WM) and cerebrospinal fluid (CSF) regions. First, anatomical images were segmented to create tissue masks for gray matter, WM, and CSF. A noise mask was then generated by combining the WM and CSF masks. The physiological noise signals were extracted from the noise mask by applying it to the fMRI time series. Principal Component Analysis was performed on these signals, and the first five principal components (PCs) were retained as noise regressors (Behzadi et al., 2007)

#### fMRI first-level GLM analysis

Analyses were conducted within the general linear model (GLM) framework in SPM to examine momentary BOLD signal modulation as a function of boundary proximity. A single condition regressor was included for TRs with translation events: when the participant was moving within the virtual environment to indicate target location, modeled with a fixed duration of 1.5 seconds to match the TR. Specifically, each translation TR was modeled as a separate event, allowing us to capture time-resolved changes in neural activity during navigation. Because participants were moving continuously during each TR, mean distance for the nearest x - and y-boundaries was computed across the full 1.5-second window. Finally, we computed the sum of these average distances to the nearest x-and y-boundaries for each translation TR event and incorporated this measure as a parametric modulator (pmod) in the GLM. The parametric modulator was included as a first-order polynomial without orthogonalization, ensuring independent estimation of the main condition and distance effects. To account for motion and physiological noise, six motion parameters (translations in x/y/z dimension, yaw/pitch/roll rotation) from realignment and five aCompCor noise components derived from PCA of the WM/CSF noise mask were included as nuisance regressors. This approach minimizes motion-related and physiological confounds, improving the specificity of neural activation estimates. A 128s high-pass filter was applied to remove low-frequency drift. Analyses were conducted in native space without spatial normalization to avoid spatial distortions or interpolation errors.

For statistical analysis, individual beta estimates for the parametric modulator (pmod) were extracted from the bilateral EC and subiculum, the primary regions of interest. To ensure that group differences in BOLD activity were not driven by differences in fMRI data quality, we quantified temporal signal-to-noise ratio (tSNR) within each ROI by dividing the mean signal intensity by its standard deviation over time. Additionally, to evaluate whether head motion during scanning contributed to group differences, we calculated each participant’s average linear and angular displacement per scan volume. Motion parameters were extracted during the realignment step in SPM12, and displacements were computed by summing translations across the x, y, and z axes, and rotations across yaw, pitch, and roll. These values were then averaged across the time series to obtain per-scan motion estimates. These measures were then used in statistical analyses to assess group differences.

Finally, to control for differences in the number of translations across age groups, we performed separate control analysis in which younger adults’ data were truncated by 19.84% to match the length of older adults’ data, and the pmod analysis was re-run. This ensured that observed effects were not driven by differences in the number of translation TR events.

### 2.6 Statistical analysis

All statistical analyses were performed in R using R studio (2023.06.2). Behavioural data analysis only included data from the fMRI scanning session (after participants were trained to a criterion). To analyze the effect of age and target location on performance and navigation behaviour, we used linear mixed-effects models as implemented in the lme4 package (Bates et al., 2012) in R. All models included age group (young vs. older adults) and target location (trash bin, plant, ball) as fixed effects, along with their interaction. Participant was included as a random effect with a random slope for target location, allowing for individual variability in target-related effects. Age group was coded using sum contrasts. To compare target locations, we used successive difference contrasts, where: the first contrast compared the trash bin vs. plant. The second contrast compared the plant vs. ball. Before analysis, we identified and removed outliers using the interquartile range (IQR) method applied to each participant’s Euclidean error distribution: we calculated the 25th (Q1) and 75th (Q3) percentiles, defined the IQR as Q3L–LQ1, and excluded any observation lying more than 1.5L×LIQR below Q1 or above Q3. Removal of individual outlier responses based on Euclidean error resulted in 1.75% of data for young adults and 2.65% for older adults being removed. This procedure was implemented to exclude occasional trials in which participants may have temporarily lost concentration or become disoriented, leading to atypical responses and navigation behaviors that deviated from their typical performance.

## 3. Results

### 3.1. Behavioral results

#### 3.1.1. Memory for object locations in older adults declines with increasing distance from boundaries

First, we examined the precision of spatial memory in young and older adults, using Euclidean distance (vm) as a measure of accuracy. Euclidean error was calculated as the straight-line distance between the participant’s recalled object location and its actual location in the virtual environment (Figure 1b). Overall, we did not find group differences between young and older adults in the accuracy of their spatial memory as indexed by Euclidean error (β = 0.61, SE = 0.37, t = 1.63, p = 0.104, Figure 2a), presumably as a result of the extensive training procedure before fMRI scanning. To ensure that the absence of an age difference in Euclidean error was not driven by speed–accuracy trade-off, we examined the relationship between response time and Euclidean error. No significant correlations were observed in either group (Figure S1), suggesting that response speed did not influence accuracy.

**Figure 2.**
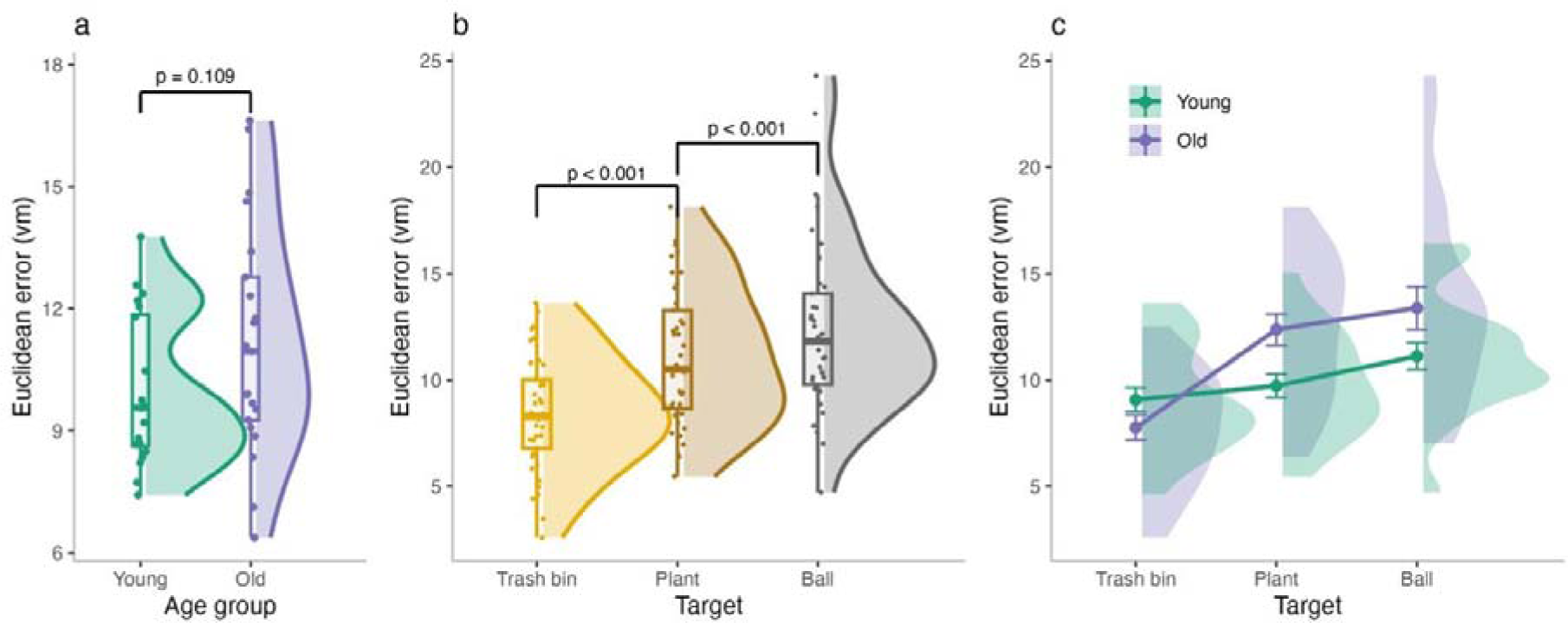
Age-related differences in spatial memory precision as a function of boundary proximity. (a) Mean Euclidean error in younger and older adults. (b) Overall Euclidean error across age groups. (c) Euclidean error as a function of target object location, with objects positioned at increasing distances from environmental boundaries. (d) Euclidean error for each target location separated by age group.

However, spatial memory accuracy was strongly influenced by the target’s position in the environment. Both groups’ error increased for the plant when compared to the trash bin, which was located closest to the boundary (β = 2.17, SE = 0.29, t = 7.38, p < 0.001, Figure. 2b). Similarly, errors increased across both groups for the ball, the object farthest from the boundary, (β = 1.69, SE = 0.37, t = 4.62, p < 0.001, Figure. 2b), compared to the plant.

Importantly, compared to younger adults, older adults’ error showed a larger increase in Euclidean error for the plant when compared to trash bin (β = 1.24, SE = 0.29, t = 4.21, p < 0.001, Figure. 2b), with no age group difference for the comparison of error between plant and ball (β = 0.53, SE = 0.37, t = 1.44, p = 0.150, Figure. 2b). The absence of an age-group interaction for the plant versus ball contrast may reflect the relatively small difference in their cumulative distances to boundaries (96vm [plant] vs 102vm [ball]), potentially reducing sensitivity to boundary-related effects. These results suggest that object locations further from boundaries are recalled with lower accuracy (c.f. Lee et al., 2018), and that this effect is more pronounced in older adults, suggesting an increased reliance on boundary information in aging.

#### 3.1.2. Older adults’ object location memory is biased towards boundaries

Visual inspection of individual responses across target locations suggested that object location estimates varied with boundary proximity in both age groups, and that these effects differed across target locations—indicating differential sensitivity to boundary proximity (Figure 3a). To systematically assess this, we calculated the boundary bias score (Figure 1b), which quantifies the ratio between the participants’ responses relative to the environmental boundaries and the correct object location. Specifically, we computed the shortest distance from each object’s correct location and the participant’s response location to the nearest walls along the x- and y-axes. Bias ratios were derived by dividing the distance to the wall for the correct position by the distance to the wall of the participant’s response, then averaging across both dimensions (x and y) to obtain a total boundary bias score. A value of 1 indicates no systematic bias, values greater than 1 reflect a tendency to place objects closer to boundaries, and values less than 1 indicate a bias away from boundaries.

**Figure 3.**
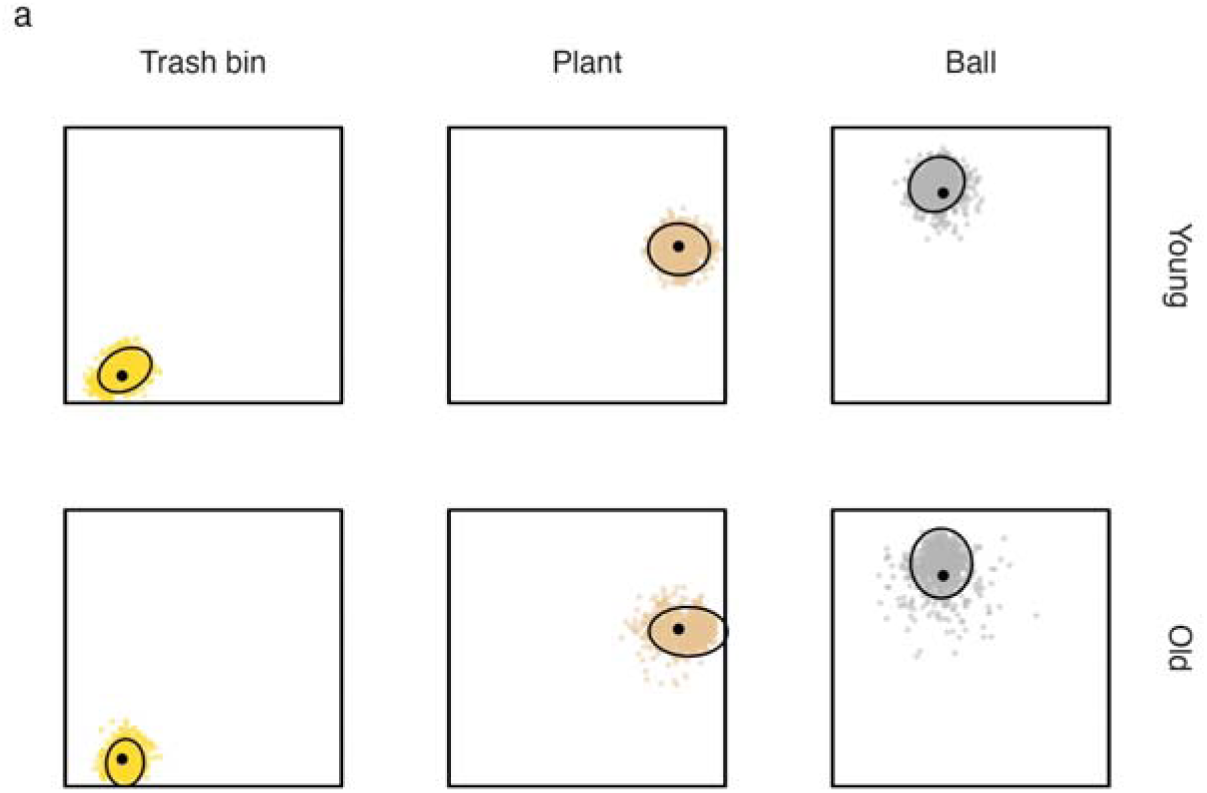
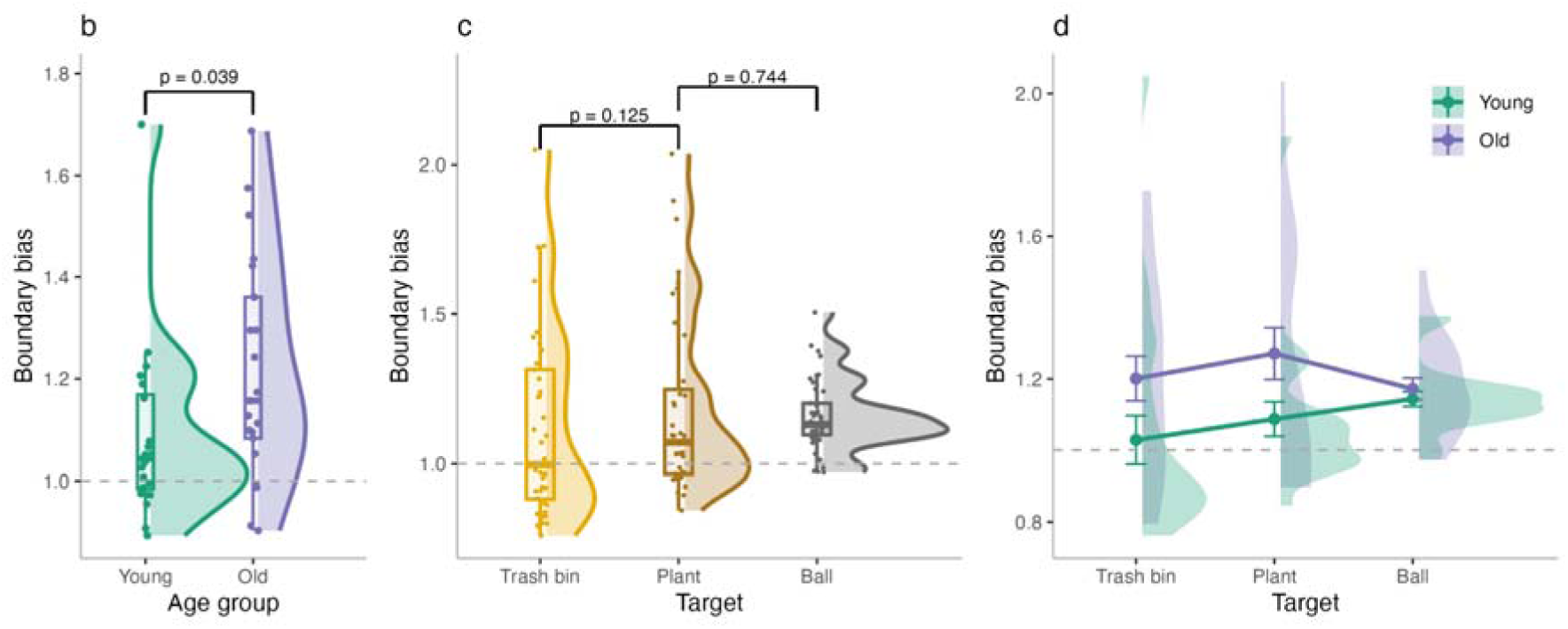
Boundary bias in younger and older adults. (a) Individual responses by target location. (b) Boundary bias scores across age groups. (c) Boundary bias scores across targets. (d) Boundary bias scores for each target separated by age group. A score of 1 indicates no systematic bias; values >1 reflect a tendency to place objects closer to boundaries, and values <1 indicate a bias away from boundaries.

We found that older adults exhibited a significantly greater boundary bias compared to young adults (β = 0.06, SE = 0.03, t = 2.06, p = 0.039, Figure 3b). Across both age groups, boundary bias did not differ significantly across target locations. Bias was numerically higher for the plant compared to the trash bin, the object closest to the boundary (β = 0.04, SE = 0.02, t = 1.53, p = 0.125), but this effect was not significant. Similarly, no significant difference was observed between the plant and the ball, the farthest object from the boundary (β = 0.01, SE = 0.02, t = 0.33, p = 0.744).

Focusing on age group and target interaction, older adults demonstrated a differential effect of target location on boundary bias when compared to younger adults. Specifically, while no significant age-related differences were observed for the contrast between the trash bin and the plant (β = -0.02, SE = 0.02, t = -0.95, p = 0.344), older adults showed a lower boundary bias increase for the ball compared to plant than younger adults (β = -0.05, SE = 0.02, t = - 2.27, p = 0.023). This effect was likely driven by a steeper increase in boundary bias across target locations in younger adults (Figure 3c), who showed relatively low or even negative bias for the trash bin (the object closest to the boundary) and progressively higher bias for more distant objects. In contrast, older adults consistently placed objects closer to boundaries across all targets, resulting in a flatter boundary bias gradient. Importantly, a supplementary analysis using distance to the closest wall rather than the combined x- and y-axis measure yielded a highly similar pattern of results (Table S1), indicating that our findings are robust to the specific boundary distance metric used. These findings suggest that while older adults generally show a stronger overall boundary bias, younger adults’ use of boundary proximity is more sensitive to target distance.

#### 3.1.3. Targets near boundaries elicit boundary-directed navigation in aging

Next, we aimed to assess navigation behavior to the response location, independent of the accuracy of participants’ spatial memory, to determine whether they tended to navigate closer to environmental boundaries than an ideal trajectory. To quantify this, we computed the boundary proximity index (BPI, Figure 1b), which measures deviations from an optimal (beeline) path in terms of boundary proximity with the endpoint of that trajectory at the response location. To compute the BPI, we first calculated the median distance to the nearest x- and y-boundaries along both the ideal (beeline) trajectory and the actual trajectory taken by the participant. BPI was then calculated by dividing the beeline distance to the response location by the actual trajectory to the response for each dimension. The resulting ratios were averaged across the x- and y-axes to produce a single BPI score. Values greater than 1 suggest a tendency to navigate closer to boundaries, while values less than 1 indicate a preference for more central navigation. To ensure robustness of this measure, we also compared BPI values derived from the mean versus the median distances and found them to be highly correlated (R = 0.94, p < 0.001; Figure S2), confirming that our results are not dependent on the choice of central tendency measure.

There were no significant differences in BPI between young and older adults (β = -0.001, SE = 0.002, t = -0.37, p = 0.712), suggesting that both groups exhibited similar overall tendencies in boundary-based navigation. However, there was substantially greater variability in BPI among older adults (SD = 0.005) compared to younger adults (SD = 0.002), as illustrated in Figure 4a. This difference in variance was statistically significant, as confirmed by Levene’s test for equality of variances (F(1, 39) = 18.19, p < 0.001).

**Figure 4.**
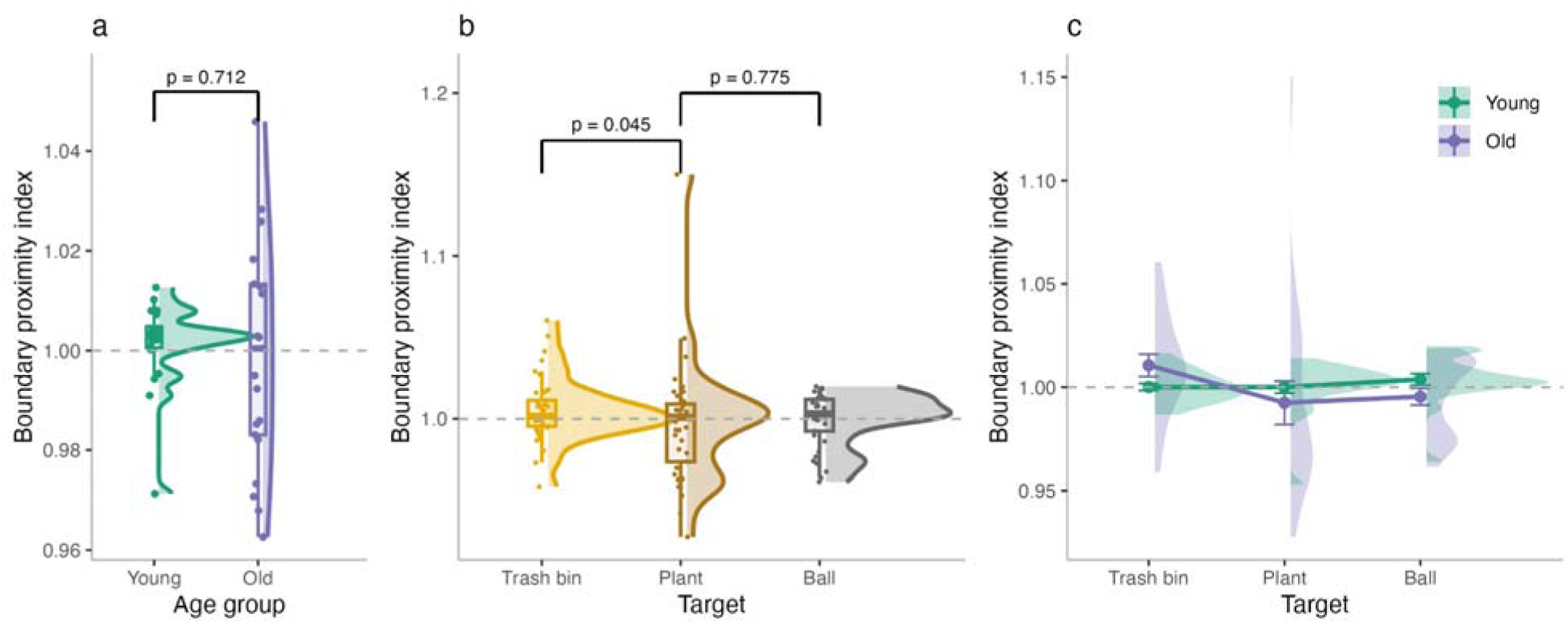
Boundary proximity index (BPI) in navigation behavior. (a) BPI across age groups. (b) BPI across target locations. (c) BPI for each target separated by age group. Values greater than 1 reflect a bias toward navigating closer to boundaries, while values less than 1 indicate a tendency toward more central navigation.

The linear mixed-effects models revealed that BPI varied as a function of target location. Participants navigated significantly closer to the boundary when heading to the trash bin compared to the plant (β = -0.005, SE = 0.002, t = -2.01, p = 0.045), whereas no difference emerged between the plant and the ball (β = -0.001, SE = 0.002, t = -0.29, p = 0.775). This pattern indicates that navigation was biased toward walls when the target was positioned near a boundary (i.e., the trash bin).

Importantly, this effect was driven by older adults, who showed a significantly larger decrease in BPI for the trash bin compared to the plant (β = -0.006, SE = 0.002, t = -2.48, p = 0.013). In contrast, no difference was observed between the plant and the ball (β = -0.003, SE = 0.002, t = -1.31, p = 0.192). Thus, interactions between age group and target location suggest that older adults tended to navigate closer to boundaries when the target was located nearby, whereas younger adults maintained relatively unbiased trajectories across all target positions (Figure 4c).

To assess whether this boundary-directed navigation observed for the trash bin in older adults was behaviorally beneficial, we correlated boundary proximity with Euclidean error separately for each target. In older adults, higher boundary proximity was associated with reduced error for the trash bin (R = –0.64, p = 0.002), whereas no such relationship emerged for younger adults or for the other targets (Figure 5). This finding suggests that older adults’ boundary-directed navigation at the trash bin location may have functionally supported their spatial memory performance.

**Figure 5.**
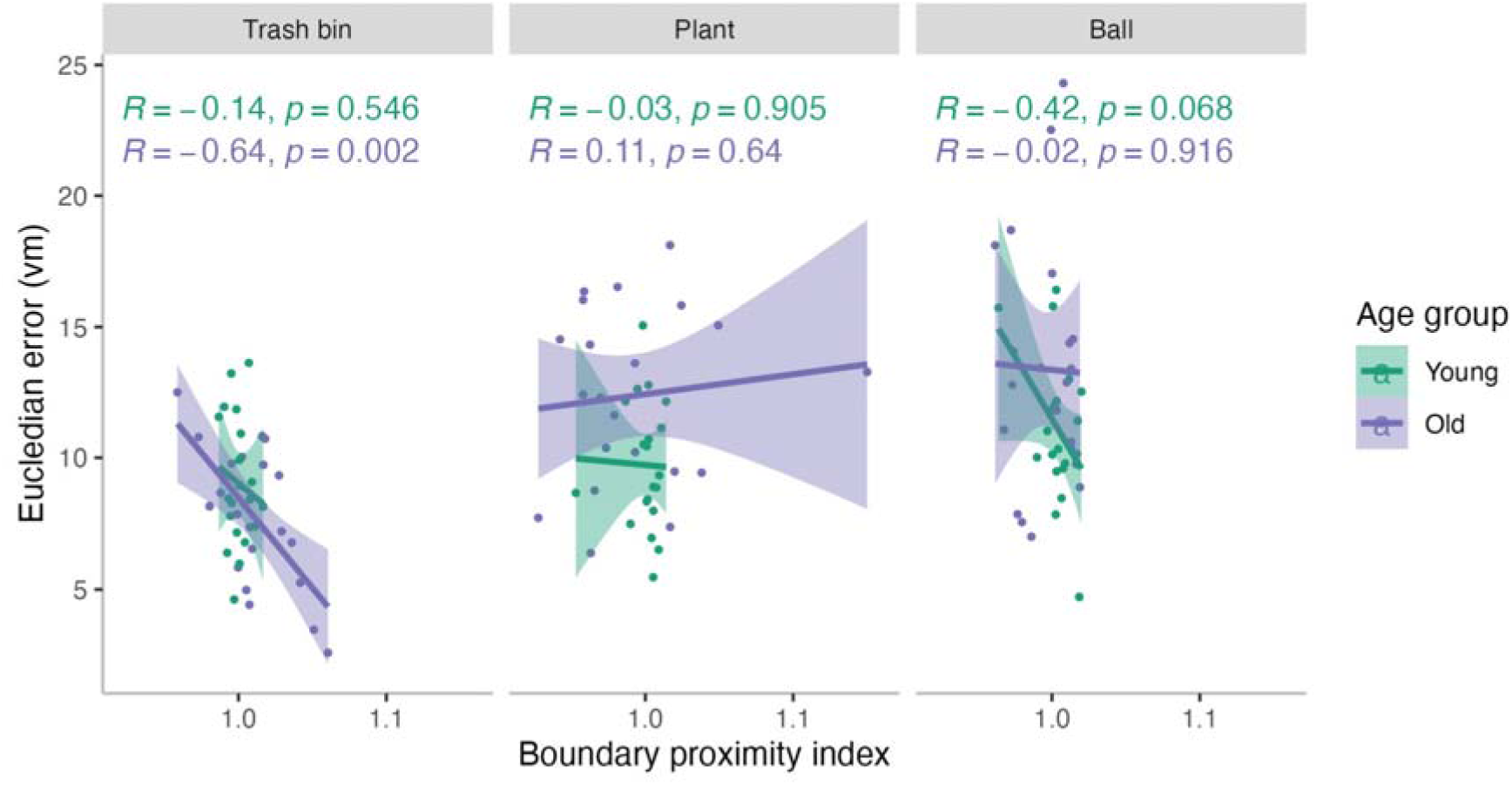
Relationship between boundary proximity and Euclidean error across target locations. Scatterplots show Euclidean error (vm) as a function of boundary proximity index (BPI) for the trash bin (left), plant (middle), and ball (right) separately for younger (green) and older (purple) adults. Shaded areas represent 95% confidence intervals of the regression lines.

In addition to metrics focusing on use of boundaries we also assessed the potential reliance on view-based strategies using path similarity index (PSI; for more information see Figure S3). Older adults showed higher PSI values for more distant targets, consistent with a shift toward view-based navigation (Figure S3a), but importantly this effect was not related to memory accuracy, suggesting that such a shift did not reliably support performance (S3b).

### 3.2. fMRI results

The behavioral findings presented above suggest that boundary processing is altered in aging, with older adults showing increased reliance on boundary proximity. To address if older adults’ representations of boundaries at the neural level are also altered, we examined moment-to-moment processing of distance to boundaries during active navigation using fMRI, focusing on the EC and subiculum—regions known to support boundary-based spatial coding. This allowed us to ask not only whether these regions track boundary distance differently in older versus young adults, but also whether these neural signals help explain the behavioral patterns observed, such as the increased boundary bias and reduced spatial precision for distant targets. In doing so, we aimed to gain a more precise understanding of how the neural representation of boundary information is affected by aging, and how such changes may give rise to the altered spatial strategies observed at the behavioral level.

#### 3.2.1 Momentary BOLD activity modulation as a function of distance to boundaries in EC and subiculum

First, we examined whether BOLD activity was significantly modulated by boundary distance in each group by testing whether the parametric modulation (pmod) values differed from zero. As all data were normally distributed (Shapiro-Wilk test, p > 0.05), we conducted one-sample t-tests separately for each group (young, older) and each region of interest (ROI; EC and subiculum). For younger adults, the modulation was not significantly different from zero in either the EC (p = 0.160, M = -0.0001) or the subiculum (p = 0.125, M = -0.001), suggesting no clear boundary-related modulation in these regions. In contrast, older adults showed significant negative modulation in both EC (p < 0.001, M = -0.004) and subiculum (p < 0.001, M = -0.003), indicating that in older adults, BOLD activity in these regions decreased significantly as distance from the boundary increased. To further evaluate the absence of significant effects in younger adults, we turned to a Bayesian framework to assess whether the data supported the null hypothesis of no boundary-related modulation. Bayesian one-sample t-tests (BF_01_) based on the Lee and Wagenmakers (2014) Bayes factor interpretation guidelines revealed anecdotal (<3) evidence for the null hypothesis in both the EC (BF_01_ = 1.72) and subiculum (BF_01_ = 1.44), suggesting weak support for the absence of modulation in these regions. We also performed a control analysis in which the younger adults’ data were truncated to match the session length of the older adult group. The pattern of results remained consistent. In the EC, Bayesian evidence also anecdotally favored the null (BF_01_= 2.39). However, in the subiculum, there was anecdotal evidence for the alternative hypothesis (BF_₁₀_ = 1.31 [BF_₀₁_ =0.76]), indicating a weak trend toward boundary-related modulation that does not constitute strong support for a reliable effect. It is worth noting that, as shown in Figure S4, younger adults spent less time near the environmental boundaries than older adults. This reduced sampling of the boundary region may contribute to the absence of significant boundary-related modulation in the younger group, particularly as border cells are known to fire primarily when subjects are very close to the boundary (Bjerknes et al., 2014; Solstad et al., 2008). However, this interpretation should be considered with caution, and further work is needed to disentangle the relationship between border cell activity, proximity to boundaries and BOLD modulation in these regions.

Next, we examined group differences in boundary-related BOLD modulation (pmod) in the EC and subiculum using linear regression. To account for potential confounds, models included motion parameters (linear and angular displacement) and tSNR in the respective region. In the EC, there was a significant main effect of age group, with older adults exhibiting more negative pmod values than younger adults (β = -0.002, SE = 0.001, *p* = 0.023, Figure 6a), indicating a steeper decline in BOLD activity with increasing distance from environmental boundaries. In contrast, no significant age group difference was found in the subiculum (β = -0.002, SE = 0.001, *p* = 0.075, Figure 6b), although the effect approached significance. The lack of age differences is likely due to a subtle shift in the younger adults’ distribution toward negative pmod values, though not to a degree that reached statistical significance. To ensure that differences were not driven by unequal sampling (i.e., number of translation TR events), we repeated the analysis using truncated data from younger adults to match the duration of the older group. The pattern of results remained consistent. In the EC, the age effect persisted (β = -0.002, SE = 0.0001, *p* = 0.012), confirming that the group difference was robust. In the subiculum, the age effect was in the same direction as for EC; however, as in the original analysis, it did not reach statistical significance (β = -0.001, SE = 0.001, *p* = 0.093), mirroring the results using the whole data from younger adults. For completeness, we also examined boundary-related modulation in the hippocampus (including dentate gyrus, CA1–3). Older adults showed significant negative modulation when tested against zero, whereas younger adults did not; however, no significant age-group difference was observed (see Figure S5).

**Figure 6.**
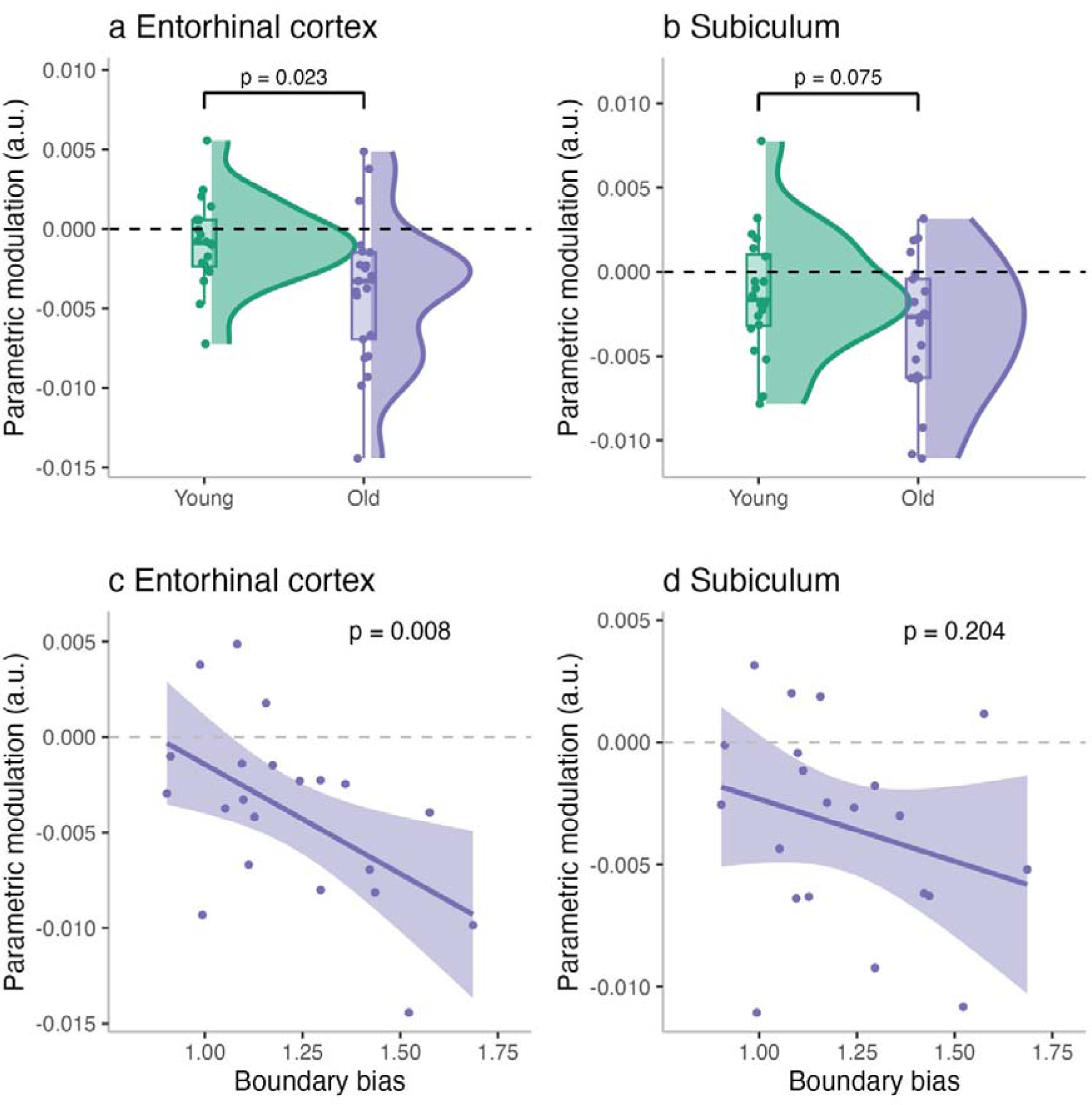
Boundary-related BOLD modulation in entorhinal cortex and subiculum. Parametric modulation (pmod) values in the entorhinal cortex (a) and subiculum (b) for younger and older adults. Negative pmod suggests decline in BOLD activity with increasing distance from boundaries (c) Scatterplot showing the relationship between boundary bias and parametric modulation in the entorhinal cortex. Stronger boundary bias was significantly associated with a more negative BOLD modulation (d) Scatterplot showing the relationship between boundary bias and pmod in the subiculum. Although the trend was in the same direction, the relationship did not reach statistical significance.

#### 3.2.2 Link between boundary-related BOLD modulation and behaviour during the task

Next, we analyzed whether individual differences in behavioral metrics—namely boundary bias and boundary navigation (BPI) —predicted the strength of BOLD modulation in the EC and subiculum. Since we did not find evidence for BOLD modulation by boundary distance in younger adults, in this analysis we focus only on older adults. We also included age and MoCA scores as covariates. In the EC, boundary bias significantly predicted BOLD modulation, with higher boundary bias associated with stronger negative pmod values (β = - 0.016, SE = 0.005, *t* = -3.00, p = 0.008, Figure 6c). This suggests that older adults who more strongly biased their memory responses toward boundaries also showed greater neural sensitivity to boundary proximity in the EC. Other predictors in the model were not significant (BPI: p = 0.826; age: p = 0.227; MoCA: p = 0.446). In the subiculum, no predictors reached significance, though boundary bias showed a numerical trend in the expected direction (β = - 0.008, p = 0.204, Figure 6d), and all other covariates were non-significant (BPI: p = 0.968; age: p = 0.755; MoCA: p = 0.346). These results highlight a robust link between behavioral boundary bias and entorhinal BOLD sensitivity to boundary proximity in older adults, while the subiculum showed a similar, though weaker, trend.

### 3.3 Spatial memory precision, task behaviour and BOLD response to boundary distance

Having examined boundary processing from both behavioral and neural perspectives, next we aimed to understand how these metrics relate to individual differences in spatial memory performance among older adults. Specifically, we asked whether boundary bias, BPI, and BOLD modulation in the EC and subiculum are associated with reduced spatial precision. To explore this, we split older adults into high- and low-performing subgroups based on a median split of participants’ mean Euclidean error and compared the behavioral and neural metrics between these groups.

Older adults with lower spatial memory precision (i.e., higher Euclidean error) exhibited significantly greater boundary bias (*M* = 1.32) compared to high-performing individuals (*M* = 1.12), *p* = 0.037 (Figure 7a). They also showed stronger negative boundary-related BOLD modulation in the EC (*M* = -0.006 vs. -0.002; *p* = 0.011, Figure 7c), with a similar trend in the subiculum (*M* = -0.005 vs. -0.002; *p* = 0.081, Figure 7d), suggesting increased neural sensitivity to proximity to environmental boundaries in lower-performing individuals. No significant difference was observed in BPI between the groups (*M* = 0.99 vs. 1.00; *p* = 0.420, Figure 7b), indicating that overall navigation tendency relative to the boundary was not reliably associated with performance. These findings suggest that stronger reliance on boundary cues, both behaviorally and neurally, may reflect a compensatory mechanism in older adults with reduced representational fidelity. Specifically, when the fine-grained precision of spatial representations declines, anchoring responses to salient environmental structures such as boundaries may provide a fallback strategy that supports task performance, albeit in a less flexible manner. Consistent with this interpretation, supplementary analyses comparing younger adults with high- and low-performing older adults (Figure S6) revealed that high-performing older adults did not differ significantly from younger adults in spatial accuracy, whereas low performers showed significantly larger errors.

**Figure 7.**
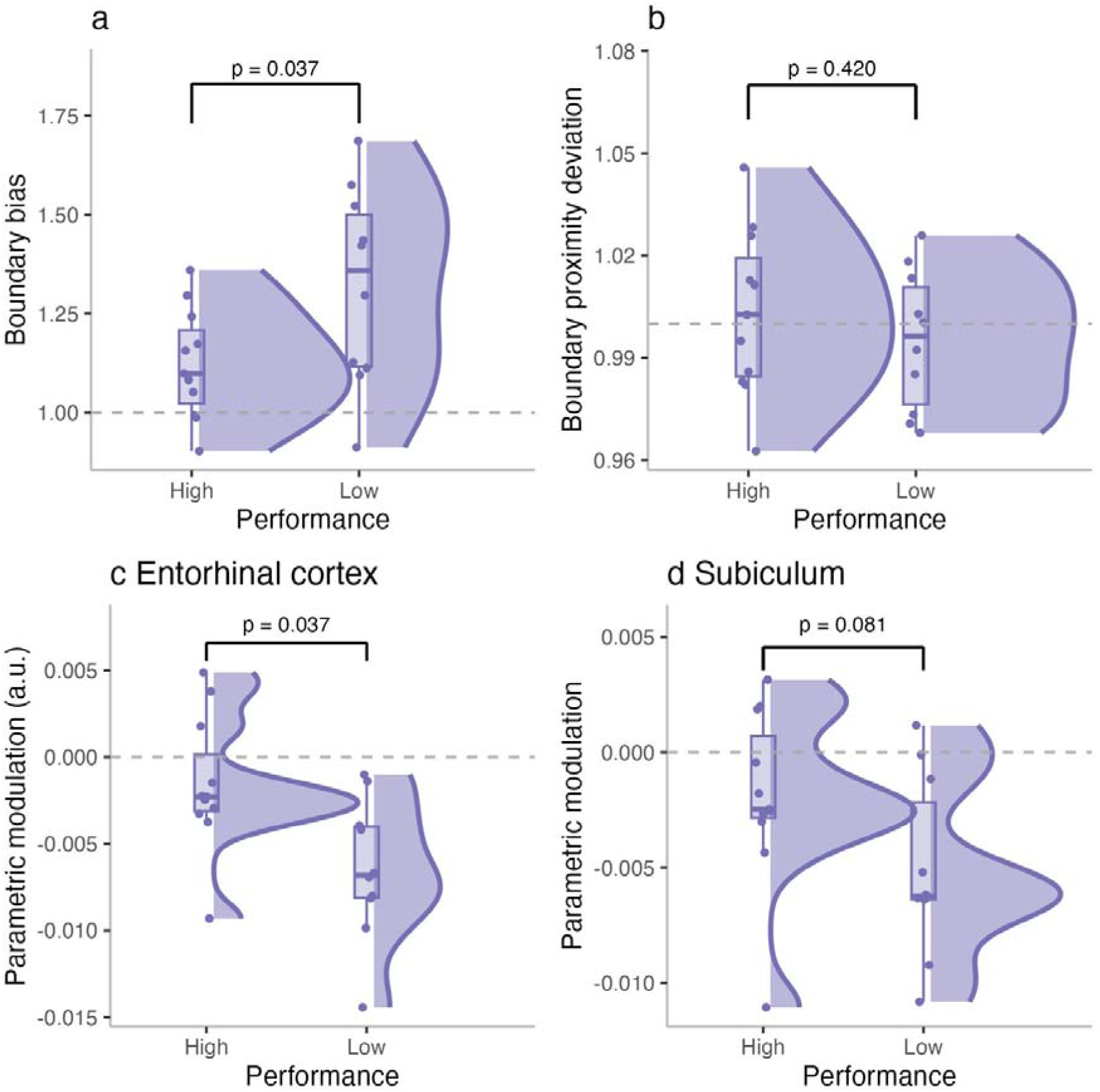
Associations between spatial memory performance and BOLD sensitivity in older adults. (a) Boundary bias scores for high- and low-performing older adults, showing significantly greater bias in the low-performing group. (b) Boundary Proximity Deviation did not significantly differ between groups, (c) Parametric modulation in the entorhinal cortex, with low performers showing significantly stronger negative BOLD modulation by boundary distance. (d) Parametric modulation in the subiculum, showing a trend towards boundary-related neural modulation in low performers.

## 4 Discussion

In this study, we investigated how aging affects the use and neural representation of environmental boundaries during an object-location memory task. While overall spatial memory precision, as measured by Euclidean error, did not significantly differ between younger and older adults, older adults showed heightened sensitivity to target location. Specifically, their memory performance declined more steeply for objects located farther from environmental boundaries, suggesting a stronger dependence of their spatial memory on boundary proximity. Furthermore, older adults showed greater boundary bias in object location memory together with boundary-related modulation of BOLD activity in the EC and subiculum. Finally, individual differences in spatial memory precision among older adults were linked to both behavioral and neural markers of boundary processing: low-performing individuals showed stronger boundary bias and more pronounced BOLD modulation by boundary distance. These findings suggest that boundary-related cues may become more prominent anchors of spatial memory in aging, particularly when precise spatial representations begin to deteriorate.

Although all participants were trained to criterion before performing the task in the scanner, age-related differences in spatial memory still emerged, but they only became apparent when performance was examined as a function of object distance from boundaries. In line with previous work (e.g., Joue et al., 2024; Lee et al., 2018), we replicated the general finding that objects positioned near boundaries are remembered with greater accuracy. Notably, this effect was amplified in older adults, who exhibited a small but reliable reduction in memory precision for targets farther from boundaries compared to younger adults. These results suggest that proximity to boundaries plays an increasingly important role in supporting spatial memory with age.

One possible explanation is that spatial representations become more uncertain or degraded with increasing distance from boundaries, particularly in older adults. Boundaries serve as stable and salient environmental anchors that help constrain spatial encoding (Brunec et al., 2018; O’Keefe & Speakman, 1987). When objects are located near these anchors, their positions can be encoded more precisely. In contrast, as distance from boundaries increases, the spatial representations of target locations likely become fuzzier, more ambiguous, and less anchored to reliable environmental structure. For older adults—whose ability to form or maintain fine-grained spatial representations may already be compromised (Diersch et al., 2021; McAvan et al., 2021; Segen et al., 2021)—this increased representational uncertainty can lead to a sharper drop in memory precision that we observed for the target locations further from the boundary. The findings highlight that while boundaries support spatial memory in general (Joue et al., 2024; Lee et al., 2018; O’Keefe & Speakman, 1987), their role becomes disproportionately important in aging, effectively compensating for declining spatial fidelity when available, but also leading to reduced performance when those environmental boundaries are less informative.

This increased dependence on boundaries may also help explain the systematic bias in older adults’ memory for object locations toward environmental boundaries. While any boundary bias differing from 1 implies greater Euclidean error, the *direction* of bias is informative: a bias < 1 would reflect systematic displacement *away* from the boundary, whereas the consistent bias *toward* boundaries observed in older adults—especially low performers—suggests strategic anchoring rather than random error or avoidance. This interpretation is consistent with the category adjustment model (Huttenlocher et al., 2000), which posits that spatial memories are reconstructed from a combination of fine-grained metric information and coarse categorical knowledge (e.g., “near the wall” vs. “in the center”). When metric precision is low—as is often the case in aging (McAvan et al., 2021; Moffat et al., 2001; Nilakantan et al., 2018; Pertzov et al., 2015; Segen et al., 2021)— responses are increasingly biased toward the center of the relevant category, or toward salient environmental boundaries. Given that environmental boundaries serve as especially reliable spatial cues (Bécu et al., 2020), they likely served as strong categorical anchors in our task, drawing responses toward them under conditions of uncertainty.

When examining the tendency to navigate closer to boundaries, group differences emerged as a function of target location. Specifically, older adults exhibited a small yet reliable preference to stay closer to boundaries for the trash bin, the object located nearest to the wall. Importantly, this tendency was associated with preserved memory precision for this target, suggesting that boundary-proximal navigation in older adults reflects a behavioral adjustment that benefits performance rather than a maladaptive bias. In contrast, younger adults’ navigation behaviour remained unbiased towards either the boundaries or the center of the environment, likely because they could maintain accuracy without relying on boundary proximity. While this increased use of boundaries in older adults could, in principle, reflect an anxiety- or safety-driven response—akin to thigmotaxis in rodents—this seems unlikely to be the sole explanation, as navigation behaviour across both groups was goal-directed and modulated by target locations i.e., navigation closer to boundaries only for the target closer to the boundary but not for the other targets. Nevertheless, it remains possible that older adults, particularly those with poorer navigation abilities, experience heightened spatial anxiety, which may contribute to altered use of boundaries during navigation. Future studies should disentangle strategic versus affective influences on boundary proximity by incorporating direct measures of spatial anxiety or arousal.

Overall, our behavioral findings suggest that older adults rely more heavily on environmental boundaries to support spatial memory, as reflected in increased boundary bias and steeper declines in accuracy with greater distance from boundaries. To probe the underlying mechanisms, we analyzed fMRI data collected during active, self-guided navigation in the EC and subiculum—regions central to boundary-based spatial coding (Bjerknes et al., 2014; Solstad et al., 2008; Lever et al., 2009; Doeller et al., 2008; Lee et al., 2018; Shine et al., 2019; Stangl et al., 2021). In older adults, BOLD activity in both regions decreased systematically with increasing distance from boundaries, whereas in younger adults, signals remained relatively stable. This pattern suggests that in aging there is a shift to stronger reliance on signals near boundaries and a steeper decline once farther away potentially reflecting a shift from flexible spatial mapping to proximity-based or cue-driven strategies.

Previous studies in younger adults have reported boundary-related activity across regions of the medial temporal lobe (Stangl et al., 2021), including the EC and subiculum (Lee et al., 2018; Shine et al., 2019). However, some findings suggest a more prominent role for the subiculum. For example, Lee et al. (2018) observed greater theta power during the encoding of locations near environmental boundaries compared to those farther away, but this effect was present only in the subiculum and not in the EC. Our results, on the other hand, revealed stronger BOLD modulation by boundary distance in the EC in older adults, with no evidence for such modulation in either the EC or subiculum in younger adults. One possible explanation for this discrepancy lies in the spatial reference frames targeted by each study. Lee et al. examined allocentric spatial relationships, specifically the distance between the target location and the boundary, and found subicular responses under those conditions. In contrast, our task focused on the distance between the participant and the boundary during movement, allowing us to capture dynamic, self-referenced distance to boundaries encoding. These two frames of reference may engage partially overlapping but distinct spatial coding mechanisms, potentially explaining the differing patterns of regional activation.

What may be driving this shift in strategy in older adults? A potential mechanism may lie in changes to the neural architecture of BVCs. According to computational models and experimental work (e.g.,Bicanski & Burgess, 2018, 2020; Hartley et al., 2000; Muessig et al., 2024), there are more BVCs closer to environmental boundaries and possess narrower tuning fields (Lever et al., 2009), whereas BVCs that encode locations farther from boundaries are fewer and exhibit broader tuning curves. If aging leads to a general degradation in BVC function—due to neural or synaptic loss (e.g. Barnes, 1988; Burke & Barnes, 2006; Du et al., 2003; Henstridge et al., 2019)—this degradation is unlikely to selectively target BVCs based on their preferred firing distance. Nonetheless, due to the inherently sparse representation of space farther from boundaries (Muessig et al., 2024), even uniform neuronal loss would disproportionately reduce spatial coverage at central locations. Since there are fewer BVCs farther from boundaries, each cell must cover a broader spatial area, meaning that loss or disruption of a single BVC results in a disproportionately larger spatial region left unrepresented or represented inaccurately. In contrast, densely-packed boundary-proximal BVCs would maintain reliable spatial coverage despite equivalent neuronal loss. This mechanism can explain the steeper decline in boundary-related BOLD modulation we observed in older adults for locations farther from environmental edges. Importantly, since some BVCs are still likely to represent locations farther from boundaries, this may account for the weaker and less consistent negative modulation of BOLD activity in the subiculum in older adults compared to EC.

In contrast, the EC is traditionally considered to contain border cells with a much more limited spatial range (Bjerknes et al., 2014; Solstad et al., 2008), making it less dependent on more widely distributed BVCs that may degrade with age. In the EC we observed more consistent and reliable boundary-related modulation in older adults. This likely reflects increased reliance on border cells, which fire only in close proximity to environmental boundaries. Importantly, border cells appear to remain functionally intact with age and even in Alzheimer’s disease models—Fu et al. (2017) reported preserved border cell firing rates in both young and older animals. Since border cells provide reliable, short-range inputs, older adults may come to rely disproportionately on these signals when near boundaries. This over-recruitment of EC border cell activity could explain the stronger boundary-related modulation we observed, but also the steeper decline in signal once participants move away from the boundary—reflecting the limited spatial range of these cells. In this sense, boundary-based navigation in aging may reflect reliance on spatial signals that remain accessible when broader spatial coding is degraded.

Notably, boundary-related navigation behavior has also been observed in individuals at genetic risk for Alzheimer’s disease (e.g., Kunz et al., 2015; Coughlan et al. 2020). For example, Coughlan et al. (2020) showed that APOE ε4 carriers navigated closer to environmental boundaries in virtual environments. This pattern may similarly be supported by preserved border cell inputs, which—while reliable—have a limited spatial range (Bjerknes et al., 2014; Solstad et al., 2008). Such a reliance may locally benefit performance when targets are near walls but can distort memory for locations farther away. This is consistent with our findings that older adults maintained comparable precision to younger adults for the object located closest to the boundary, while showing larger errors for more distant targets. At the same time, older adults also demonstrated a tendency to navigate closer to the wall for the nearest object (trash bin), suggesting that boundary-proximal cues provided reliable support when available, but at the cost of reduced precision and increased bias when targets were located further from boundaries. Together, these patterns suggest that altered boundary processing in aging may reflect both attenuated contributions of long-range BVCs in the subiculum and increased reliance on border cell input in the EC, a division that helps explain the differential neural results across regions and their behavioral consequences.

This framework also offers a plausible explanation for the behavioral boundary bias we observed. If BVCs further from boundaries are not only fewer in number (Muessig et al., 2024) but also more broadly tuned (Bicanski & Burgess, 2018; Lever et al., 2009), then age-related degradation could cause a shift in the center of mass of neural firing toward cells with closer boundary preferences. This would result in a biased spatial signal, effectively skewing the memory trace closer to the boundary than the true object location. Consequently, older adults’ responses would increasingly reflect locations anchored near the boundary, not because of intentional strategy, but due to biased or degraded neural representations. This mechanism complements our interpretation that boundary bias in aging arises from reduced spatial fidelity and increased reliance on structurally-defined cues when precise spatial coding fails.

This interpretation also aligns with models proposing that path integration and boundary signals provide complementary inputs to the hippocampal–entorhinal system (Evans et al., 2015). Path integration, supported by grid and head direction cells, provides a continuous metric of location but is prone to cumulative error and therefore requires correction by stable environmental cues such as boundaries (Hardcastle et al., 2015). Given that aging has been associated with reduced fidelity of grid-like representations and impaired path integration (Stangl et al., 2018), boundary inputs may help stabilize spatial representations. Our finding that older adults showed stronger boundary-related modulation in the EC, together with increased boundary bias, is consistent with such a compensatory weighting toward environmental anchors when internal spatial coding becomes less precise.

Performance variability is a well-established hallmark of cognitive aging, especially in domains like spatial memory and navigation (Moffat & Resnick, 2002; Zhou et al., 2023). Consistent with this, the behavioral and neural markers of altered boundary processing observed in our study were not uniformly present across all older adults. Our results showed that low-performing older adults—those with the highest spatial memory errors—were the ones who exhibited both stronger boundary bias and steeper BOLD signal reductions in the EC and subiculum as a function of distance from boundaries. This pattern suggests that the BVC-related degradation described above may not characterize aging as a whole, but is particularly pronounced in individuals with reduced spatial fidelity. In these individuals, boundary-based coding appears to be most disrupted, leading to biased neural representations and greater reliance on proximity-based cues. The fact that stronger boundary-related neural modulation was associated with poorer performance supports the idea that these responses do not reflect effective use of boundary information, but rather a maladaptive shift driven by degraded spatial coding mechanisms. Together, these findings provide compelling evidence that altered boundary processing contributes to the spatial memory decline observed in a subset of older adults.

## Limitations and future directions

While our findings offer important insights into boundary-based spatial processing in aging, several limitations should be acknowledged. First, the task included only a limited number of target locations, each consistently paired with a specific object category. This design does not allow us to fully disentangle the effects of boundary proximity from potential influences of object identity, familiarity, or salience. For example, individual differences in memory performance or neural responses might be partly driven by preferences or prior experiences with particular objects, rather than spatial factors alone. Additionally, the relatively small differences in boundary distance between certain locations (e.g., plant vs. ball) may have limited sensitivity to detect target effects, as similar distances to boundaries can lead to comparable performance. Nevertheless, spatial precision in our task was still influenced by these manipulations, even when the differences were relatively small, however it could have precluded some of the age specific effects. To avoid both of these potential confounds, future studies should orthogonally manipulate object identity and boundary distance across a broader range of values, ideally in a between-subjects design where objects are randomly assigned to locations for each participant to prevent object effects from being directly coupled with spatial location.

Finally, we did not collect biomarker data, such as amyloid or tau status, to rule out the presence of preclinical Alzheimer’s disease pathology in our sample. It is therefore possible that the observed deficits in some older adults may reflect early neurodegenerative processes rather than healthy aging alone. In particular, it will be important for future work to test whether the subgroup of older adults who show stronger reliance on boundaries are also those with elevated AD pathology. Such studies could provide a critical link between boundary-related navigation strategies in aging and the broader literature on grid cell dysfunction and path integration impairments in Alzheimer’s disease.

## Conclusion

In summary, our findings provide converging behavioral and neural evidence that aging is associated with a shift in how boundaries are used and represented during spatial memory. Critically, the observed behavioral patterns—such as increased boundary bias and steeper declines in memory precision with distance from boundaries—appear to be a manifestation of altered boundary processing in older adults. As boundary signals become less informative at greater distances, they may no longer reliably support the formation of flexible, map-like spatial representations rooted in boundary vector cell coding. Instead, older adults increasingly rely on boundaries as static cues or beacons, which may simplify spatial decision-making but limit adaptability and precision. Importantly, this shift was most pronounced in low-performing older adults, further highlighting that the observed boundary bias is not merely a strategic choice but likely reflects a deeper disruption in boundary-based coding mechanisms. This altered processing appears to contribute directly to the spatial memory and navigation deficits observed in aging. Together, these findings suggest that disruptions in boundary-based spatial computations may be a key factor underlying age-related declines in spatial memory and navigation.

## Acknowledgments

This work was supported by the Velux Stiftung (1809).

## Notes

### Competing Interest Statement

The authors have declared no competing interest.

