## Supplementary material for "Altered coding of environmental boundaries in human aging: an fMRI study": Supplimentary materials

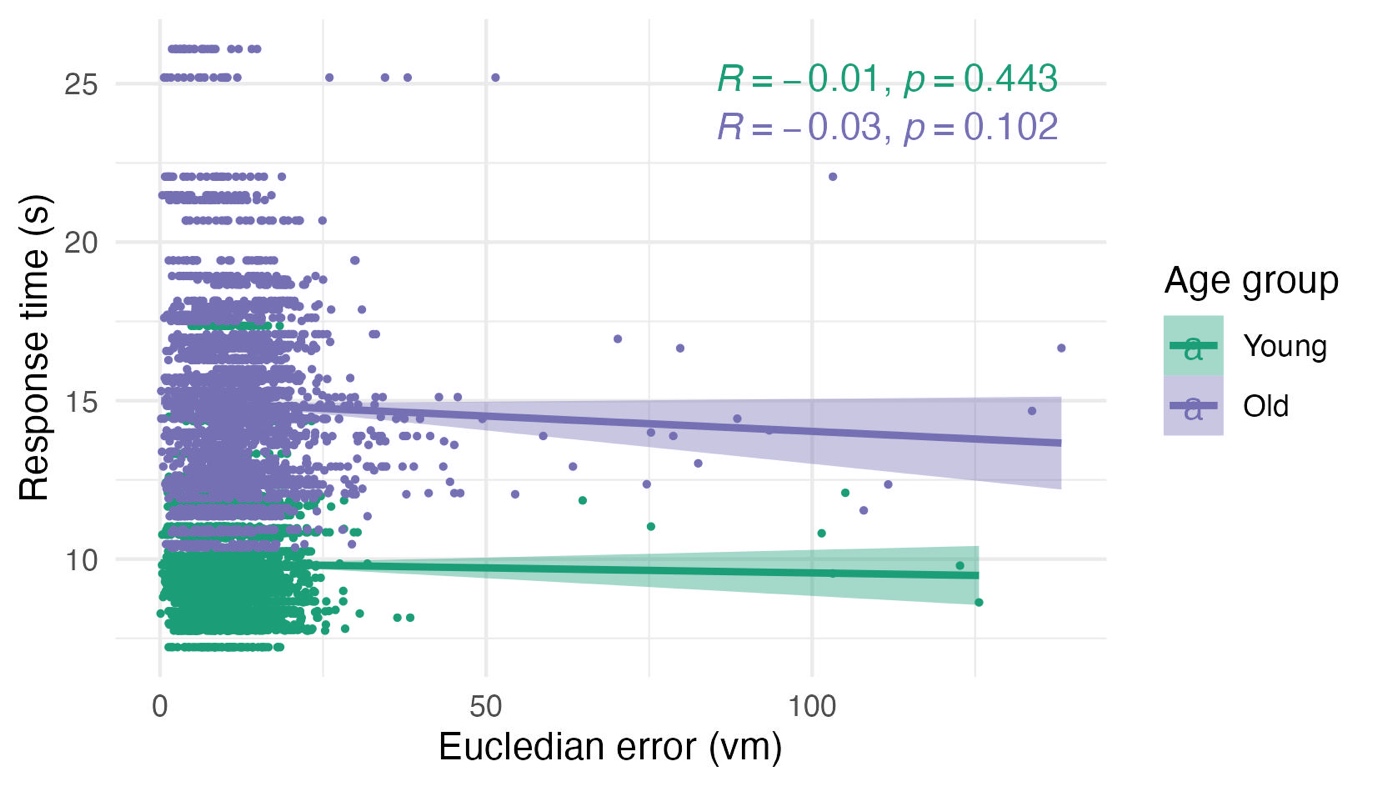

**Figure S1. Relationship between response time and Euclidean error in younger and older adults**. Each point represents the mean response time and Euclidean error for one participant, averaged across all trials. Shaded regions indicate 95% confidence intervals of the regression line. No significant correlations were observed in either age group (Young: R = –0.01, p = 0.443; Old: R = –0.03, p = 0.102).

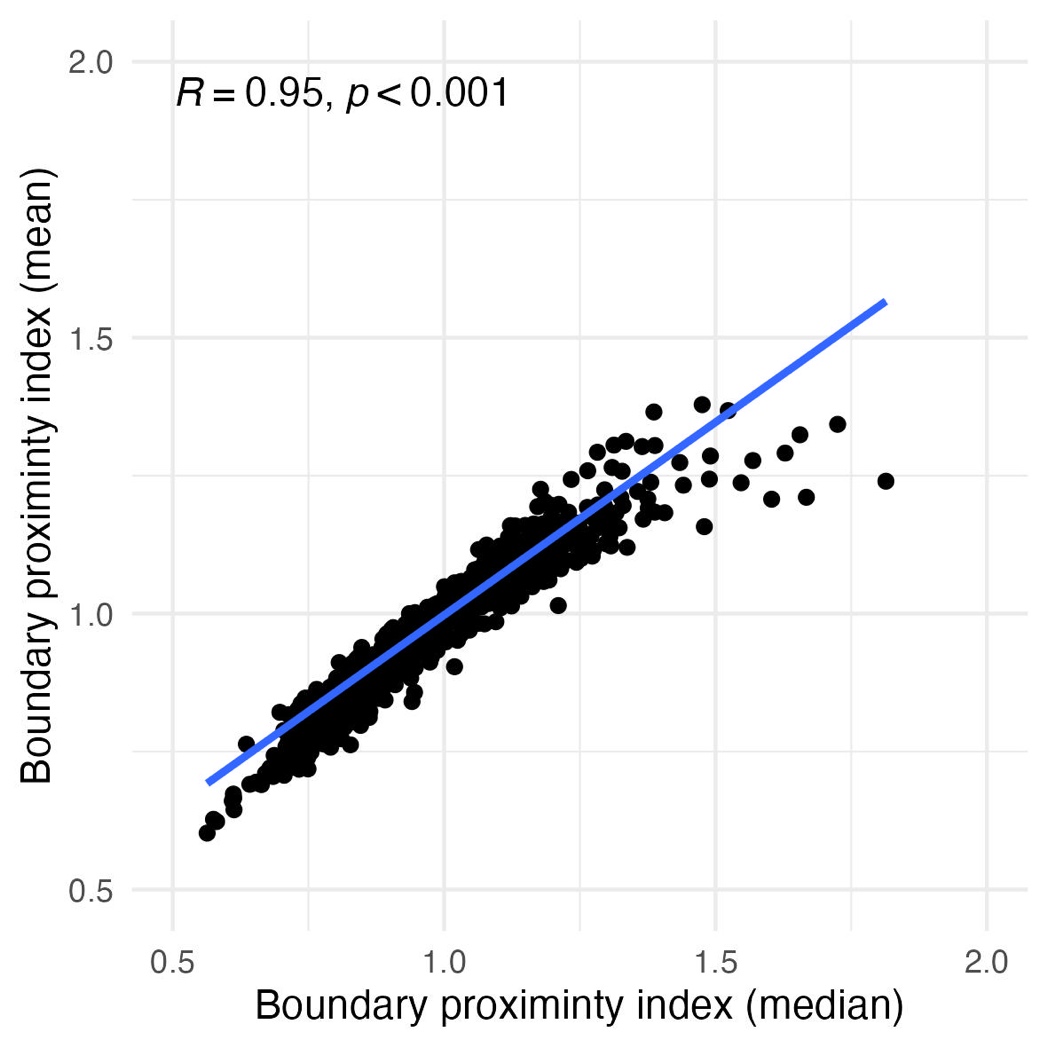

**Figure S2. Correlation between mean- and median-based boundary proximity index (BPI) values.** Each point represents an individual trial. The two measures were highly correlated (R = 0.95, p < 2.2e-16), confirming that BPI provides a robust measure of boundary-related navigation behavior regardless of whether mean or median distances were used in its calculation.

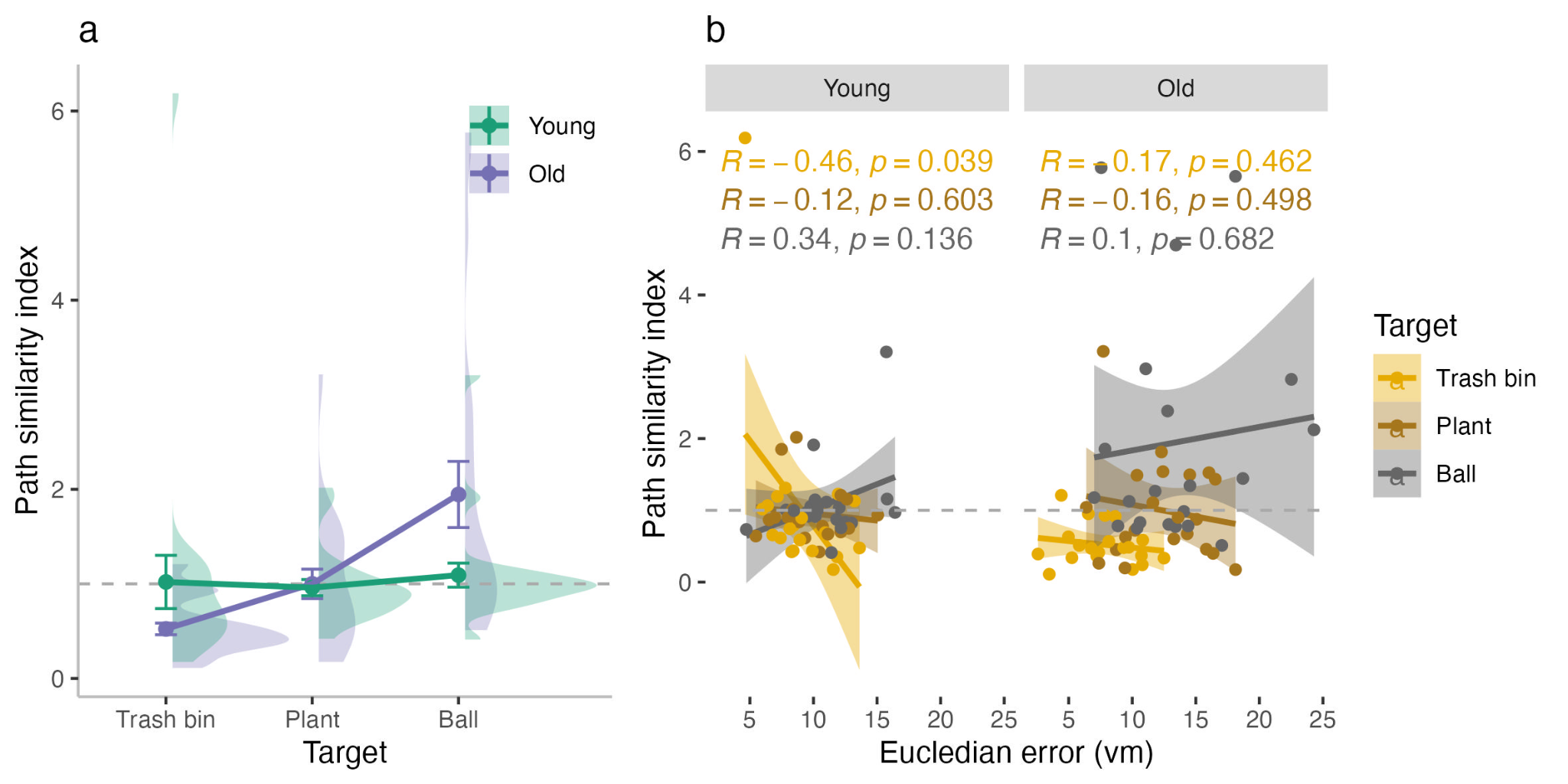

**Figure S3. Path similarity index (PSI) as a proxy for view-based navigation.**

PSI was calculated as the ratio between variance in the direct (beeline) heading to the target and variance in the actual approach heading during the final quarter of each trajectory. Values greater than 1 indicate that approach headings were more consistent (lower variance) than the direct heading—suggesting potential reliance on view-based strategies (e.g., visual snapshots). Values below 1 indicate less consistent approach headings. (a) PSI across targets and age groups. There were no differences between the age groups, however we did observe target effects with higher PSI values for the plant vs trash bin (β = 0.30, CI = 0.11–0.50, *p* = 0.002) and ball vs plant (β = 0.42, CI = 0.23–0.61, *p* < 0.001). These main effects of target are largely driven by older adults who show increased PSI for plant vs trashbin: β = 0.30, CI = 0.11–0.49, *p* = 0.003 and for ball vs plant: β = 0.35, CI = 0.16–0.54, *p* < 0.001. Importantly, only for the ball, were the PSI index higher than 1 in older adults (b) Correlations between PSI and Euclidean error by target and age group. Notably, for the ball—the only object where older adults showed PSI values >1—PSI was not significantly correlated with memory accuracy, suggesting that while older adults may shift toward a more view-based strategy, this shift did not reliably affect performance.

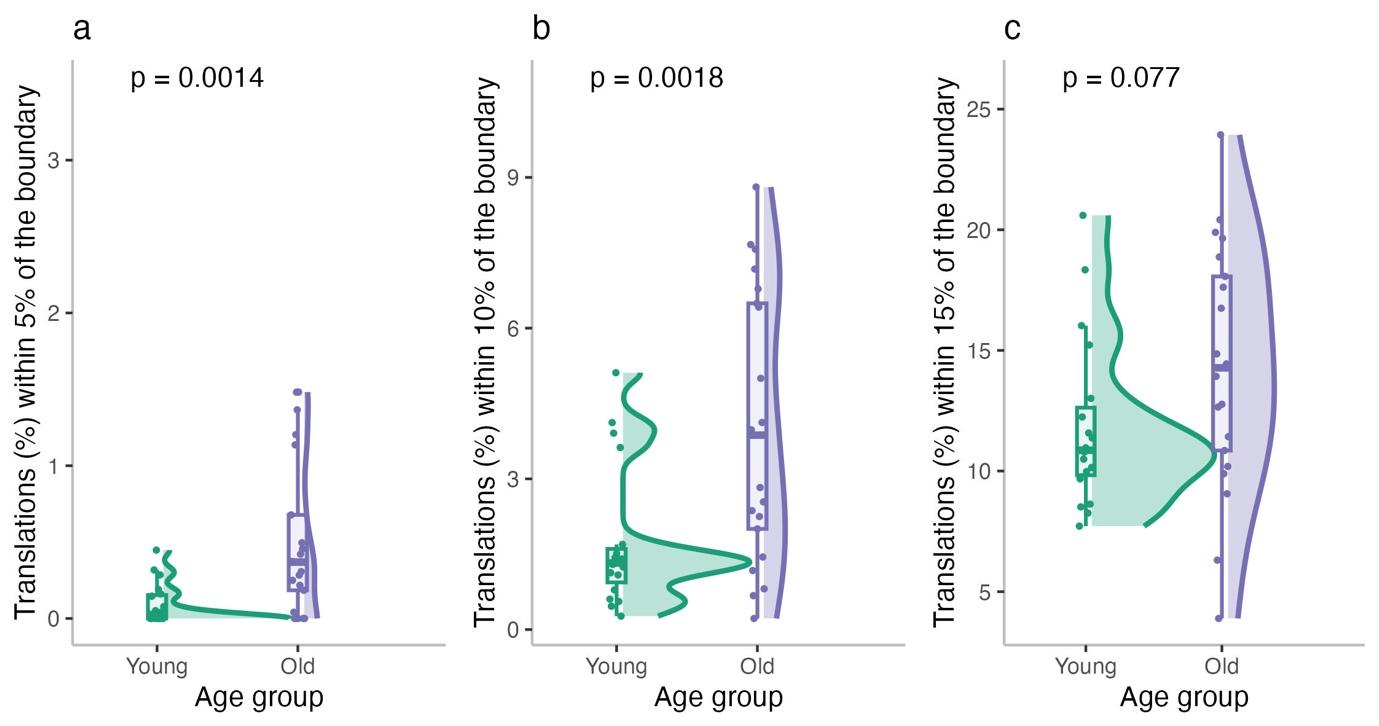

**Figure S4. Age differences in boundary proximity during translation events.**
(a–c) Percentage of translation events occurring within 5% (a), 10% (b), or 15% (c) of the environmental boundary, for young and older adults. Individual data points, group distributions (half-eye plots), and boxplots are shown. P-values are from independent-samples t-tests.
*One younger adult participant was excluded from all analyses shown here due to having values more than 3 standard deviations above the young adult group mean and was considered a statistical outlier.*

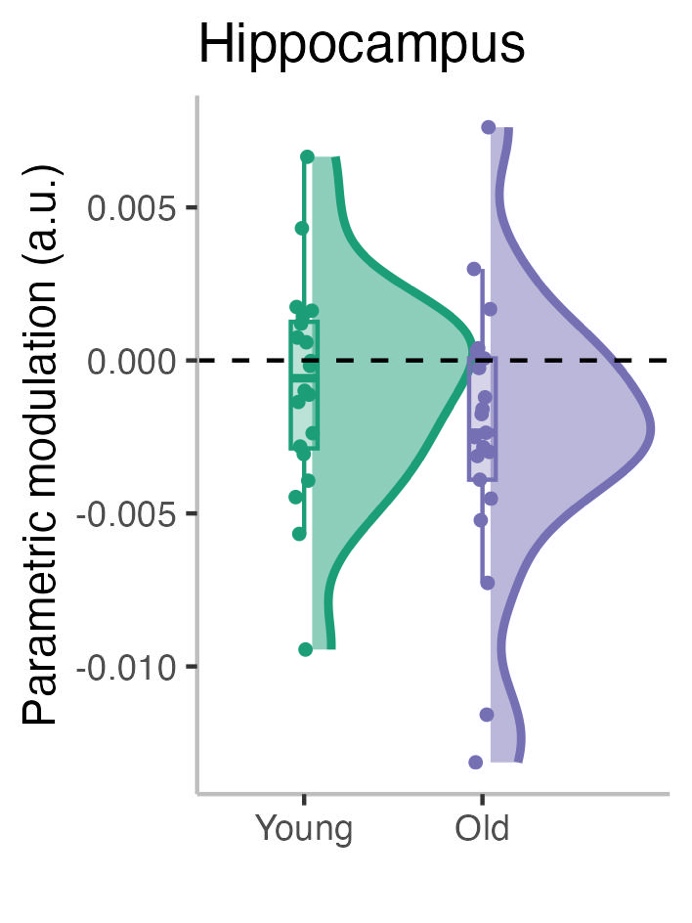

**Figure S5 Hippocampal parametric modulation results.**

Parametric modulation (pmod) by boundary distance in the hippocampus (including DG, CA1–3) for younger (green) and older (purple) adults. One-sample t-tests against zero showed a significant negative modulation in older adults (p = 0.023) but not in younger adults (p = 0.298). However, a linear model including head motion and tSNR as covariates revealed no significant group differences (β = –0.00054, t = –0.69, p = 0.494). These results suggest that while older adults show some boundary-related modulation in the hippocampus, effects are attenuated compared to the entorhinal cortex and subiculum and do not differ reliably between groups.

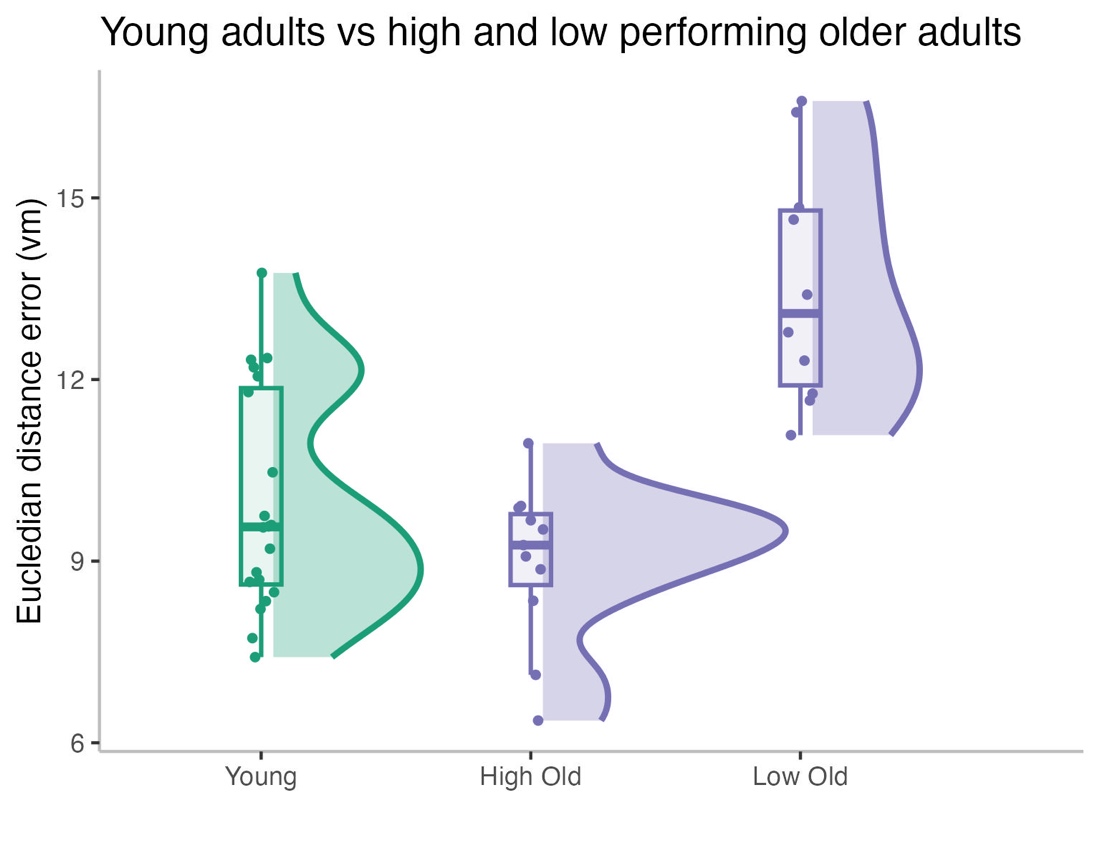

**Figure S6. Comparison of Euclidean distance error between younger adults and high- versus low-performing older adults**. Violin/box plots show the distribution of median Euclidean error per participant. High-performing older adults exhibited accuracy comparable to younger adults (M = 9.0 vs. 10.0 vm; t(29) = 1.53, p = 0.14), whereas low-performing older adults showed significantly larger errors than younger adults (M = 13.6 vs. 10.0 vm; t(28) = –4.89, p < 0.001). These results support the interpretation that boundary anchoring in low-performing older adults reflects a compensatory mechanism for reduced spatial fidelity, while high-performing older adults maintain accuracy at levels similar to younger adults, likely via more flexible strategies.

**Table 1 Comparison between Total Boundary Ratio and Closest Boundary Ratio**

| **Predictor** | **Total Boundary Ratio** | **Closest Boundary Ratio** |
| --- | --- | --- |
|  | *β* (SE), *t*, *p* | *β* (SE), *t*, *p* |
| Intercept | 1.15 (0.03), 36.90, *p* < .001 | 1.28 (0.06), 22.109, *p* < .001 |
| Age Group | **0.06 (0.03), 2.06, *p* = .039** | **0.14 (0.06), 2.44 *p* = .015** |
| Target1 | 0.04 (0.02), 1.53, *p* = .126 | 0.06 (0.04), 1.36, *p* = .172 |
| Target2 | 0.01 (0.02), 0.33, *p* = .744 | -0.02 (0.04), -0.58, *p* = .563 |
| Age Group × Target1 | -0.02 (0.02), -0.95, *p* = .344 | -0.06 (0.04), -1.44, *p* = .149 |
| Age Group × Target2 | **-0.05 (0.02), -2.27, *p* = .023** | **-0.10 (0.04), -2.33, *p* = .020** |
